# VERA: agent-based modeling transmission of antibiotic resistance between human pathogens and gut microbiota

**DOI:** 10.1101/356824

**Authors:** OE Glushchenko, NA Prianichnikov, EI Olekhnovich, AI Manolov, VE Odinzova, AB Tyaht, ES Kostrukova, EN Ilina

## Abstract

The resistance of bacterial pathogens to antibiotics is one of the most important issues of the modern health care. The human microbiota can accumulate resistance determinants and transfer them to pathogenic microflora by means of horizontal gene transfer (HGT). Thus, at the moment it is important to develop methods of prediction and monitoring of antibiotics resistance in human population. We represent the agent-based VERA model, which allows to simulate the spread of pathogen with account of possible horizontal transfer of resistance determinants from commensal microbiota community. The model considers the opportunity of residents to stay in the town or in medical institution, have wrong self-treatment, reception of several antibiotics types, transfer and accumulation of resistance determinants from a microbiota to a pathogen. In this model we have also created assessment of optimum intensity of observation of infection spread among the population. Investigating model behavior we show a number of nonlinear dependencies, including the exponential nature of dependence of total of the diseased on average resistance of a pathogen. As the model infection we considered infection with *Shigella* spp, though it could be applied to a wide range of other pathogens.

**Availability and implementation:** Source code and binaries VERA and VERA.viewer are freely available for download at github.com/lpenguin/microbiota-resistome. The code is written in Java, JavaScript and R for Linux platform.

## Introduction

Increase in mortality from pathogens, resistant to antibacterial substances, at the moment is one of the most important problems of modern health care. According to the report by Review on Antimicrobial Resistance, chaired by Jim O’Neill (O’Neill, 2016), annual mortality from infections complicated by antibiotics resistance (AR) is estimated to 700,000. By 2050 it is prognosed that mortality will reach 10 million, and economic losses - 100 trillion US dollars (O’Neill, 2016). The major factor promoting the spread of resistance is wide use of antibiotics in medicine and agriculture (Rolain, 2013; Rhouma et al.;). Such factors as misuse of antibiotics by population, insufficient control of turnover of the medicines containing antibiotics, uncontrolled use of these medicines in agriculture and others (Rhouma et al., 2016; Liu et al., 2016; Nguyen et al., 2016; Österberg et al., 2016). Accumulation of AR by human microbiota commensals and transfer of AR determinants to pathogens by means of HGT could complicate this problem further (Sommer et al., 2009).

Thus, there is a need for development of the instruments of monitoring and prediction of spread of infections to human populations taking into account above-mentioned factors. It is important to use modern methods of functional genomics, next-generation sequencing (NGS) and modern computational methods, including computer modeling and also their combination.

Mathematical and computer models help to investigate and prognose the behaviour of various complex systems including systems of biological nature (Eubank, et al., 2004; Pitman et al., 2012). They are were already applied for studying the spread of AR (Spicknall et al., 2013; Fofana et al., 2017; Weinstein et al., 2001). Models exists which consider the problem of AR pathogens in hospitals (D’Agata et al., 2007; Webb et al., 2005; Austin et al., 1999). Considering the use of several antibiotics (Cohen et al., 2004). The majority of them is based on the use of differential equations (Lipsitch et. al., 2000; Colizza et al., 2007; Kaufman et al., 2009; Spicknall, et al., 2013; van Bunnik and Woolhouse, 2017). However, it was noted that agent based modeling is more suited to model infectious diseases (Willem, et al, 2015). We didn't aware of any existing models which takes into account accumulation and transmission of AR determinants in microbiota.

We offer the tool - the agent-based model VERA, which simulates accumulation and spread of pathogens resistance among the population. It can be useful for the assessment of an epidemiological state both in some place (i.e. a city, town, village etc) or in the various institutions (for example, kindergarten). Agents can be placed nominamally in the town or in the hospital, and in various states: healthy, with disease in a incubation period, with disease in an active phase. Agents microbiota accumulates resistance determinants, transfer them to the pathogen which could be transferred to other agents. Optimal monitoring intensity is evaluated based on minimization of cost functional. The result of simulation is detailed log of state transitions. Also a number of statistics could be visualized in a pdf report or displayed through the local WEB page.

## Materials and methods

### Model description

Agents are people who can be localized in a town or in a hospital. Agent can catch pathogen and change its state from healthy to state of disease in incubation period (See Fig. Schema). After the end ot the incubation period agent transits to the disease state with resistant or sensitive pathogen until it is cured either in home or in a hospital. While curing at a home it takes Antibiotic-1 either correctly or in non optimal regime (i.e. preliminary reducing the dose or cessation of treatment). Wrong home treatment makes pathogen resistant to Antibiotic-1. Patients are treated with the other antibiotic (Antibiotic-2) in the hospital. We assume that there is no formation of Antibiotic-2 resistance. Agents can be hospitalized because of the reason other then considered pathogen, and they can be infected there.

### Model of microbiota resistance formation and its transfer to pathogen

Pathogen can be in one of the two states: resistant to one of the antibiotics (Antibiotic-1) or sensitive to it. Each agent in our model has its own level of microbiota resistance, which is increasing in process of using Antibiotic-1. Also there is a constant Antibiotic-1 resistance increase (i.e. by food consumption with subtherapeutic Antibiotic-1 doses [Moore et al., 2013; Pehrsson et al., 2016; Hoang et al., 2017]), until it reaches some constant level. It is supposed that patients without heavy diseases symptoms are treated at home with Antibiotic-1, where the AR-formation by means of horizontal genes transfer through plasmids/transposons is possible (Levy et al., 2004; von Wintersdorff et al., 2016). The patients in the hospital are supposed to be treated with the other antibiotic (Antibiotic-2). We suppose that formation of Antibiotic-2 resistance by means of plasmids / transposons transfer bearing AR-determinants is improbable. With each step in simulation microbiota of each patient taking Antibiotic-1 increases resistance by *r*_*+*_, otherwise it decreases by *r*_-_. At each step we count the average microbiota resistance level to an Antibiotic-1 *r*_*m*_ and average pathogen resistance *r*_*p*_.

The possibility that the healthy agent will be infected with a pathogen and that pathogen will become resistant corresponds to the Bernoulli random distribution magnitude with possibility of success of *p*_*2*_ and *r*_*t*_ accordingly.

It is possible that pathogen will gain resistance to Antibiotic-1 with probability *p*_7_. If the pathogen acquired resistance, probability to be hospitalized on treatment increases to *p*_5_=(*p*_3_ * *r*_c_ * 10), where *p*_3_ is probability of hospitalization with AB sensitive pathogen from home treatment and *r*_*c*_ characterize the ease of pathogen acquire resistance determinants. There is also a probability that the agent in {Non Infected person in hospital} will catch a sensitive pathogen *p*_6_, there is a random Bernoulli distribution magnitude with probability of success of *r_h_* to catch a resistant pathogen.

### Model of antibiotic resistance spread and agent transmission dependences

Agents change their states, according to probabilities, and have the variable microbiota resistance level.

Probability to catch the pathogen *p*_2_ is defined dynamically on every moment of modeling and proportional to the number of infected people in town divided to a total number of people in town (see Supplementary Table S2)

We assume that the treatment course in hospital is chosen in such a way that provides recovery of the patient and without AR formation. A modeling time unit is 1 day. In the conditions of treatment (except for the treatment in hospital) the resistance increases by *r*_+_ every day with antibiotics taking (AB) and, on the contrary, decreases when reception of AB stops, at a size *r*_-_ until it races constant level *r*_*l*_.

All probabilities of transitions have Bernoulli distribution with the appropriate probability of success (see Supplementary Table S2). Transitions from one state to another are carried out by agents on each time point of modeling, according to the following rules. Agents can be in one of a number of states which differ in location (town or hospital) and the status of infection. They are: the healthy person (Healthy person), the person who addressed to the hospital not because of infection with the considered pathogen (Non Infected person in hospital), the person in incubation period (Person in incubation period), the person who is on treatment at home with the correct period of antibiotics reception (Person in AB treatment period), the person who is on treatment at home with the reduced period of antibiotics reception (Person in AB wrong treatment period), the person infected with the current pathogen and who is on treatment in hospital (Infected person in hospital), the person in the incubatory period with a resistant pathogen after the wrong treatment (Person in incubation period 2). Transitions are possible on every moment (tiks) of modeling. The short description of the states used in modeling is given below. The workflow of the VERA is presented in Figure 1.

**Figure 1.**
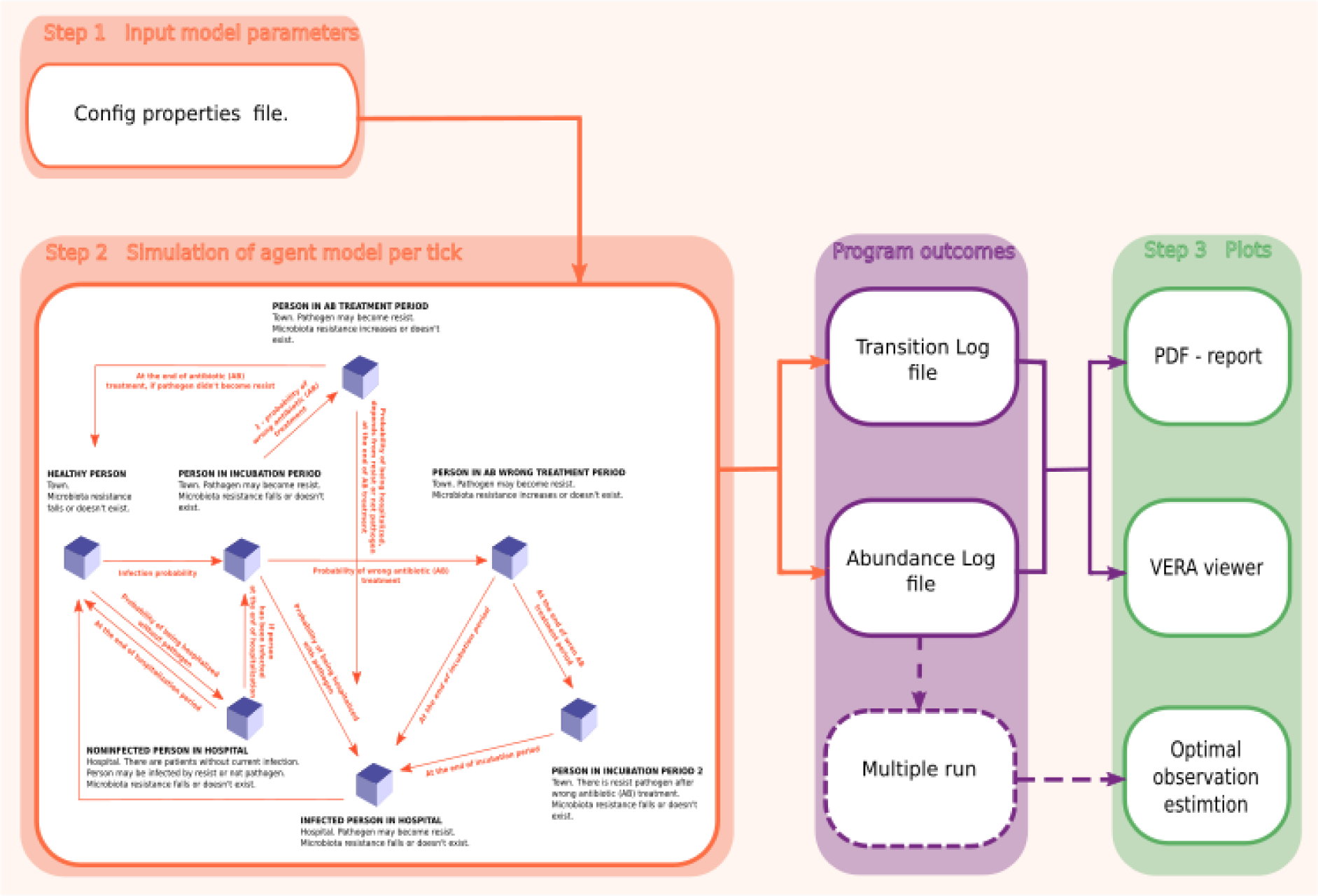
The workflow of the VERA model. Step 1 - set input parameters; step 2 - start simulation with to get: a) the table with the number of agents in each state on each time point of modeling, b) the file with transitions of each agent; step 3 - dynamic visualization, pdf-report are generated and optimal frequency of observations is calculated.

*Healthy person. State 1.* Person locate in the town without pathogen infection and reduction. The agent can get into a state *{Non Infected person in hospital}* with the possibility of *p*_1_ into *{Person in incubation period}* with possibility of *p*_2_.

*Non Infected person in hospital. State 2.* People who addressed to the hospital not because of infection with a target pathogen. Agents can go to this state only from state {Healthy person}. The intestinal microbiota AR of the person is reduced by *r*_-_, as it is supposed that there is no reception of specific antibiotic. After treatment period *i*_*p*_ agent passes back into *{Healthy person}* or with possibility of *p*_6_ is infected with a pathogen and transits to a state *{Person in incubation period}*, with probability *p*_9_ to catch resistant pathogen.

*Person in incubation period. State 3.* Agents who were infected by pathogen and with disease in the incubatory period. Pathogen can become resistant with the possibility *p*_7_. Incubation period should be defined in input parameters to match the pathogen under consideration. When incubation period is expired the person can pass into a state *{Infected person in hospital}* with possibility of *p*_3_, or be treated at home with non optimally (state *{Person in AB wrong treatment period})* with possibility of *p*_8_ or optimally (*{Person in AB treatment period}*) with probability 1 -*p*_8_.

*Person in incubation period 2. State 4.* Person gets to this state after the wrong treatment, the pathogen is supposed to be resistant. As soon as the incubation period stops, agents are hospitalized (state *{Infected person in hospital}*). Agents in hospital take other antibiotic (Antibiotic-2).

*Infected person in hospital. State 5.* Agents have the treatment in hospital. The resistance of a microbiota decreases on *r*_-_ because of change of an antibiotic to the one, where there is no resistance through horizontal genes transfer. After the treatment expiry the agent passes into a state *{Healthy person}*.

*Person in AB wrong treatment period. State 6.* In this state agents' the pathogen can become resistant *p*_7_. During the treatment at home with the possibility of *p*_4_ or *p*_5_ the agent can be hospitalized on treatment. If after the end of the treatment the agent wasn't hospitalized, he passes into a state Person in incubation period 2.

*Person in AB treatment period. State 7.* At any day of treatment agents can be also hospitalized with possibility of *p*_4_ or *p*_5_ and, also, with the treatment period expiry (in a state *{Infected person in hospital}*), if the pathogen became resistant. And if not, then with a certain level of microbiota resistance it passes into a state *{Healthy person}*.

### The optimal observation intensity estimate

We developed a procedure to estimate the necessary frequency of monitoring the infection dynamics in the town. To predict spread of infection and pathogen resistance one should monitor epidemiological state sufficiently often. But high costs of such procedures (i.e. molecular methods to determine AR determinants) requires some optimal value to be chosen to get balance between observation costs and threat of error. We implemented a pFor this, a large number of model runnings is required. So it's possible to make a multiple run of the model (see multiVERA script https://github.com/lpenguin/microbiota-resistome/blob/master/scripts/multiVERA.sh) (we used 500 tests). Let 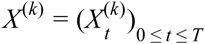 is some random process, in our case, the number of people who get sick with *k*-running (the sum of the number of infected people in the town and in the hospital, the InfectedPersonslnTown and InfectedPersonslnHospital columns in the corresponding table), where *t* - is the observation day, *T* - is the simulation time. Then consider the jump multivariant process 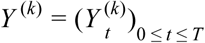, which is the prototype of the processes observations 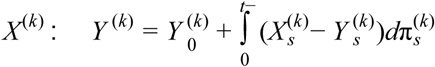 with the initial values 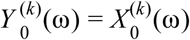. Random processes 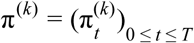 are Poisson processes with intensities λ^(*k*)^ when monitoring the process is *X*(_*K*_). Construct the estimate 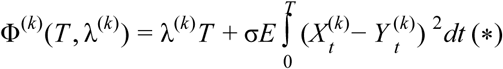. Here λ^(*k*)^*T* serves as the measurement rate, and σ is the error rate. To determine the optimal intensity of observations, it is necessary to solve the minimizing problem for the objective functional (*)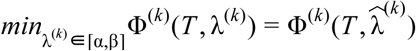, where 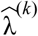 is the desired optimal observation intensity (OOI). The evaluationReportRUN_v2.sh script implements this task and determines the OOI (Supplementary Figure S1, 500 runs with 100-day simulation). By default, the error rate σ is set according to the CDC data on the number of diseases and deaths associated with AR is 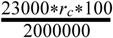.

### Parameters estimation for shigellosis

Shigellosis is widespread disease in places of public catering, day-care centres etc, consequently we investigate *Shigella* spp as a model pathogen. The value of input model parameters are by default reflected in additional materials and correspond to infection with *Shigella* spp. (see Supplementary Table S1). We consider that Antibiotic-1 is co-trimoxazole same, Antibiotic-2 is from groups of fluoroquinolone or nitrofuran.

## Results

### Qualitative analysis of the model

We got multiple launch (500 runs) of the VERA model with the same input parameters and estimated the scale of changes in the four main model indicators (the number of infected people, the probability of being infected, the average level of microbiota resistance, and the pathogen separately). This procedure allowed to assess the model results stability relative to the probability transitions. The results are shown in Figure 2.

**Figure 2.**
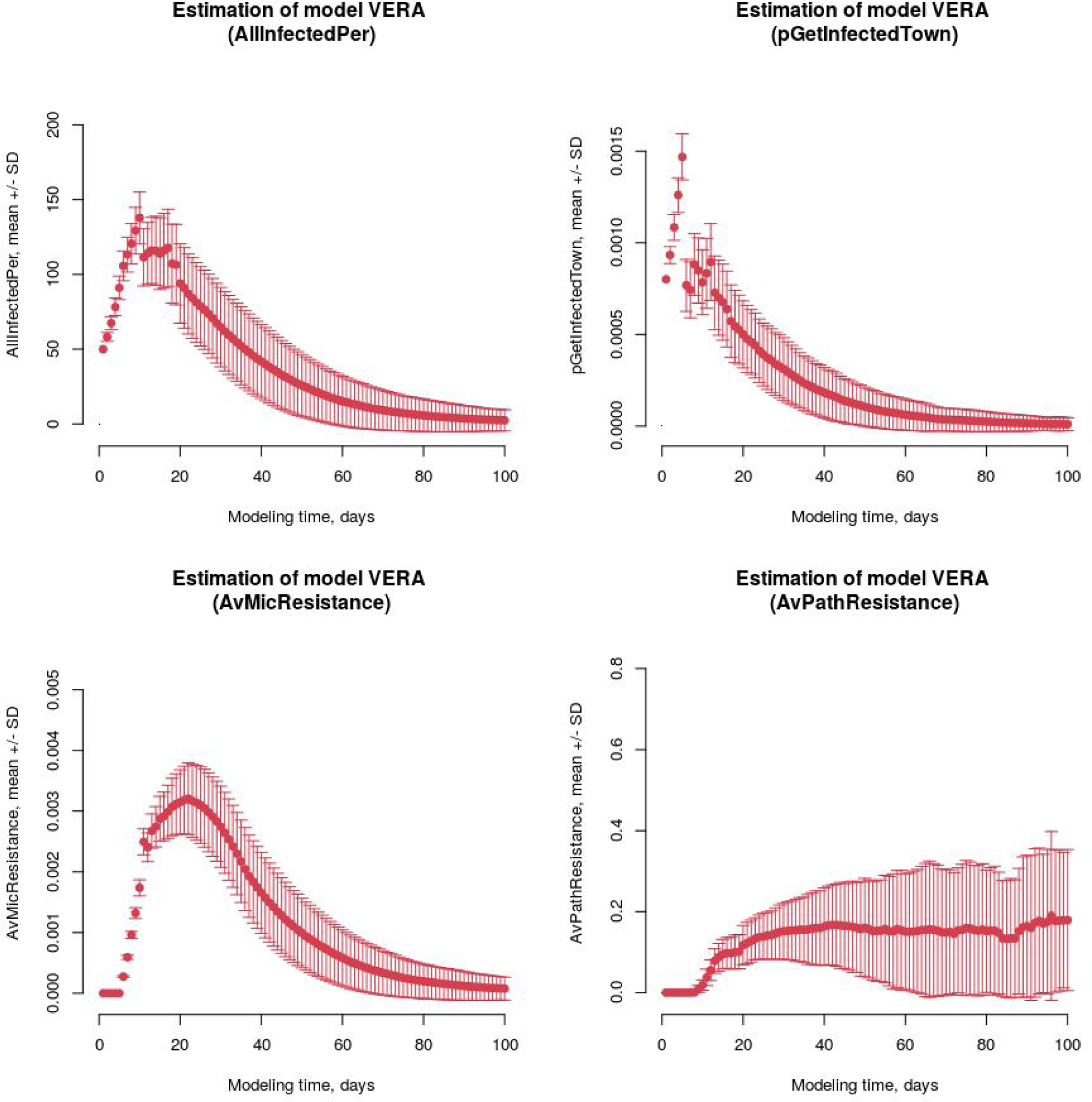
Results of the model stability assessment. The average of 500 runs relative to each simulation time point with standard deviation. a) the number of infected people (AllInfectedPer), b) the probability to be infected *p*_*2*_ (pGetlnfectedTown), c) the average level of microbiota resistance rm (AvMicResistance), d) the average level of pathogen resistance rp (AvPathResistance).

It is seen that the values of the indicators for multiple launch of the model with the same values of the input parameters differ from the mean with some deviation. The standard deviation varies with respect to the simulation time. In the case of the indicator, the number of infected people the maximum value of the standard deviation from the mean for multiple launches is explained by the data kind, here this number of people, with size of 10,000 people.

The obtained data allow us to state that this model is fairly stable and the observed spread of the values of the main parameters does not affect the change in the behavior of the model.

### CeTKa napaMeTpoB

To determine under what initial conditions there are transitions from a low infection degree to a pandemic and assumptions of the interrelation nature (or relationship type) of input parameters, a regression analysis of multiple model launches was conducted, where the simulation time was 300 days. To define the model state, we will use the values of the simulation indicators. Indicators in the VERA model are 1) the number of infected people, 2) the probability of being infected, 3) the average level of microbiota resistance, and 4) the pathogen separately. As an indicator assessment during the entire modeling period, the area under its (indicator) curve was chosen: the larger the area at a certain start-up, the higher the index under selected input parameters. To determine which of the input parameters contribute the most to changes in model values, a linear regression was constructed for each indicator with 100,000 simulations with varying input parameters (Figure 3).

**Figure 3.**
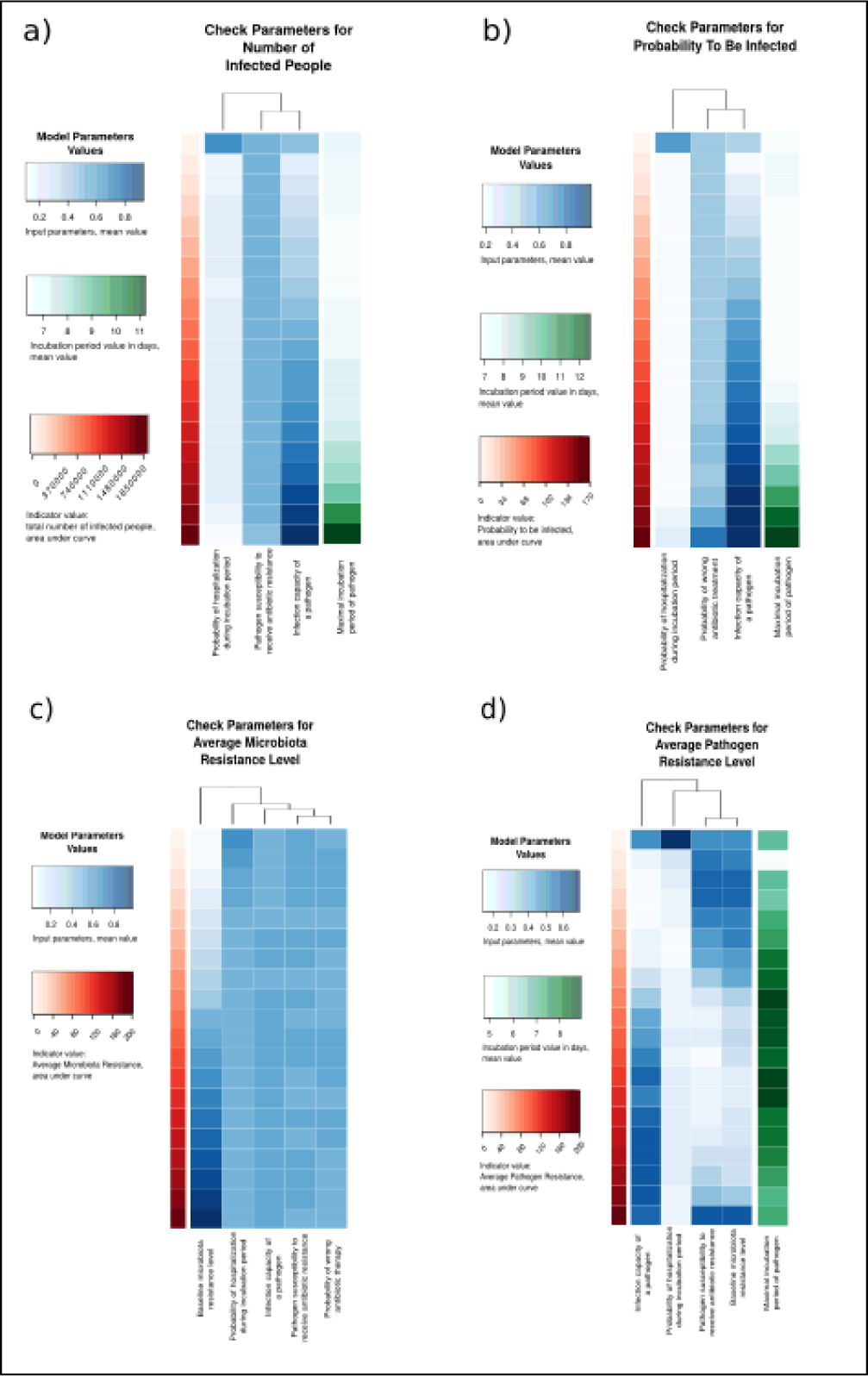
Parameter grid. Heatmap of The model indicators values depending on the control parameters values. There are the input parameters as the control parameters, which make the most significant contribution to changes in indicator values, according to regression analysis. a) The model indicator is the number of infected agents. Control parameters - the probability to be hospitalized after the incubation period, the incubation period, the pathogen susceptibility to receive antibiotic resistance (the potential of accepting an AR pathogen, how much it is susceptible to the reception of AP genes), the pathogen potential for infection. b) The indicator of the model is the probability to be infected in the town. Control parameters are: the probability to be hospitalized after the incubation period, the probability of wrong treatment, the incubation period, the pathogen potential for infection. c) The model indicator is the average level of microbiota resistance. Control parameters are: the permanent level of gut antibiotic resistance, the probability to be hospitalized after the incubation period, the pathogen susceptibility to receive antibiotic resistance (the potential of acceptance of AR by the pathogen, how much it is receptive to the reception of AP genes), the pathogen potential for infection, the probability of wrong treatment. d) The indicator of the model is the average level of pathogen resistance. Control parameters are the permanent level of gut antibiotic resistance, the probability to be hospitalized after the incubation period, the pathogen susceptibility to receive antibiotic resistance (the potential of accepting an AR pathogen, how much it is susceptible to the reception of AP genes), the pathogen potential for infection, and the incubation period.

Linear regression was constructed based on the results of 10^5^ runnings of the VERA model, where the simulation time is 300 days. To execute each such launch, config.properties file was generated with the values of the input parameters, which are randomly selected from the appropriate allowable ranges. Further, for each model indicator (the number of infected people, the probability of being infected, the average level of resistance of the microbiota and the pathogen separately), the score (the area under the indicator curve) was measured. To calculate the linear regression, the values vector for indicator was taken as the dependent variable, and the free variables are the values of the input parameters. The parameters whose contribution to the indicator change was the most significant (the value of the regression coefficients) were selected as significant for this indicator.

It turned out that the pathogen infection potential has a significant effect on the all model indicators values. Among the other parameters, the control parameters were: the incubation period *i*_*p*_, the probability of wrong antibiotic treatment *p*_8_, the pathogen susceptibility to receive antibiotic resistance *r*_*c*_ (the potential of accepting an AR pathogen, how much it is susceptible to the reception of AP genes), the permanent level of gut antibiotic resistance *r*_*l*_, the probability to be hospitalized during incubation period *p*_3_.

Thus, using a grid of parameters for model indicators, it is possible to predict the pandemic degree in the prognosis for 300 days based on the parameter values. Also a grid of parameters revealed the relation type of input parameters. The analysis has established that a permanent level of gut antibiotic resistance of agents affects only the average resistance of the microbiota and pathogen. The increased probability of wrong antibiotic treatment leads to an increased probability to be infected in the town.

### Model indicators dependencies

The main model indicator is the number of infected agents (and this value is most easily estimated in practice). Model indicators are interdependent so we tested the dependence type of pandemic state estimate on the number of infected agents and other indicators. It turned out that in the case of probability to be infected, the dependence is logarithmic (Fig. 4, a). With the increase in the probability to be infected *p*_*2*_, a rapid increase in the number of infected people and a transition to a saturation are observed. That means the saturation does not occur as quickly as its growth. The relation between the number of infected agents and the average level of pathogen resistance are characterized by an exponential dependence (Fig. 4, b). Critical values of the disease incidence are achieved only with a high proportion of resistant strains on average, and any restriction for the growth of the disease incidence does not have any noticeable effect. It can be assumed that this pattern is due to home treatment, which does not have the proper effect against resistant pathogens.

**Figure 4.**
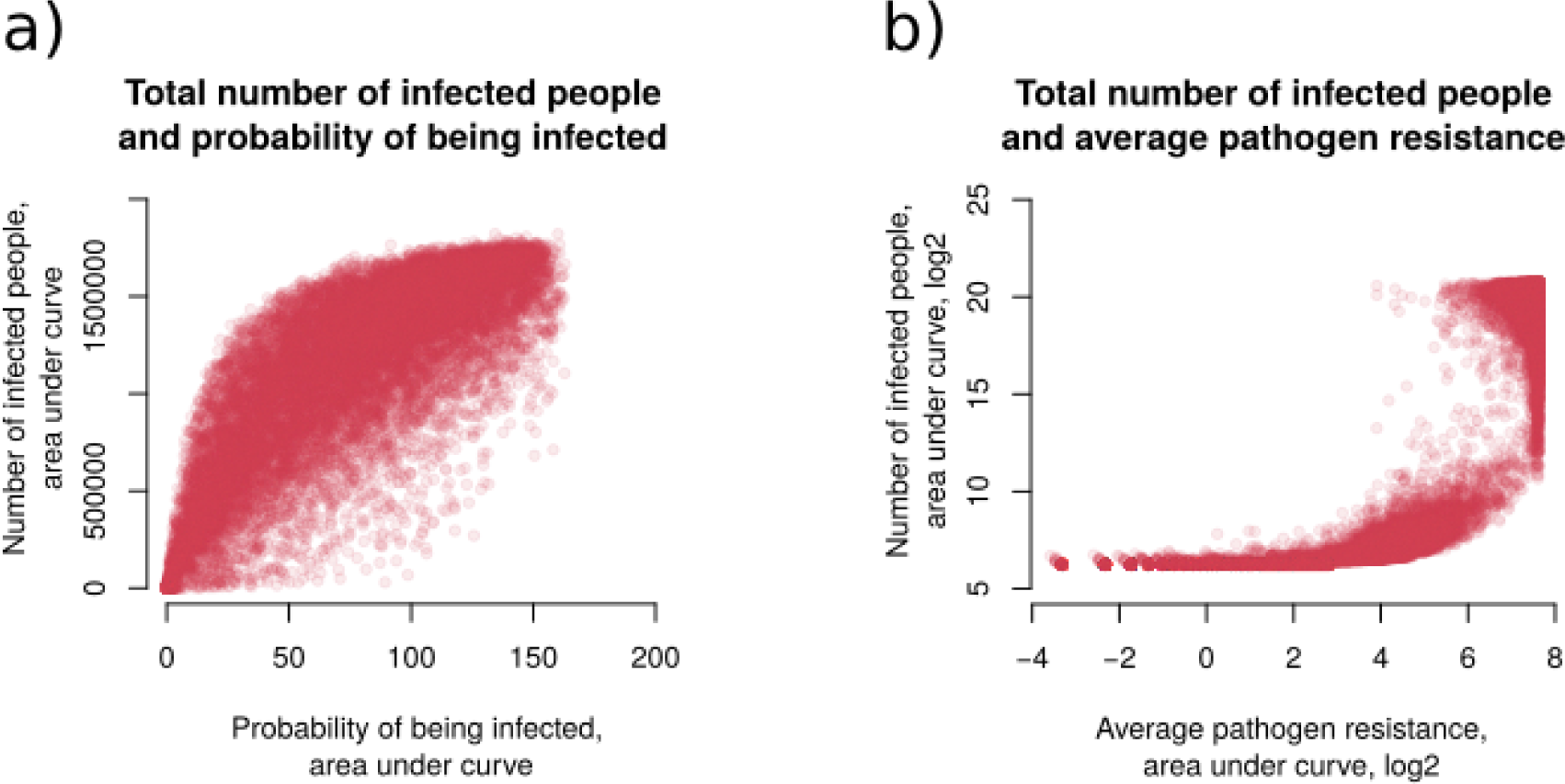
The graph (Visualization) of the infected agents number and other model indicators. a. The probability to be infected. b. The average level of pathogen resistance, along the axes of log2 values.

### Demonstration of the model using *Shigella* spp

To demonstrate the work of VERA on the model infection, we chose shigellosis (infection with *Shigella spp.*). The values of the input parameters were derived from the literature (see Supplementary Table S1). We consider that Antibiotic-1 is co-trimoxazole same, Antibiotic-2 is from groups of fluoroquinolone or nitrofuran.

In the case of *Shigella* spp. using the simulations of our model to develop an effective treatment strategy, the following recommendations can be made. It was found that the optimal observation intensity of the epidemiological situation development is two days (Figure, suppl). After multiple model run, it can be concluded that the pandemic is falling by median for 80 days (Fig. 5). Also seen is the second possible scenario, when during the development of infection we have two peaks of a pandemic. For example, as seen from one of the simulations (Fig. 5), the first peak is observed on day 65, the second at 167 after a fall of 134 days. In this case, the number of cases was close to zero at 236 days. This behavior is explained by the increased number of people who chose the wrong treatment (R = 0.865, the Spearman's rank correlation coefficient) and as a result of an increase in the average level of resistance of the microbiota and pathogen. At the time of the second peak, the resistance of the pathogen begins to reach the full saturation level. This explains that the number of infected agents begins to decline, so as only treatment in the hospital will be effective.

**Figure 5.**
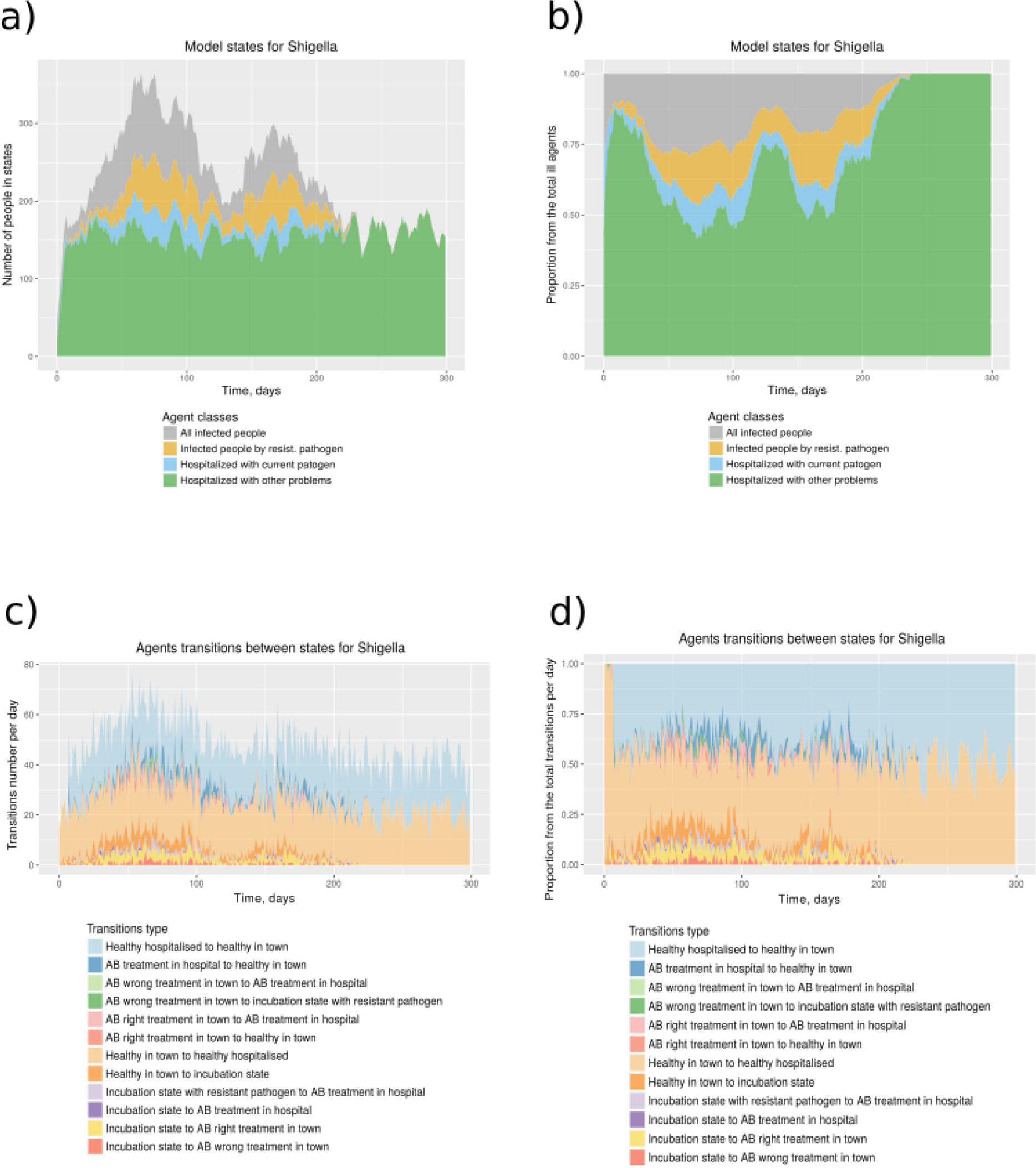
**a)** The number of agents that have been infected with the pathogen or are on treatment at each simulation moment (day). Gray indicates the total number of infected agents, of which the resistant pathogen-infected agents (yellow), hospitalized in the hospital with the pathogen (blue). Also, the number of agents hospitalized with non-pathogen infection (green color) is indicated. b) The proportion of agents that have been infected with the pathogen or are on treatment at each simulation moment (day). c) The number of agents transitions from one state to another at each simulation moment (day). d) The proportion of agent transitions from one state to another at each simulation time (day).

## Discussion

The spread of the bacterial determinants of resistance to antibiotics occurred from a long time (D'Costa et al., 2011, Perron et al., 2015) and it is not in itself a threat. However, our intensive use of antibiotics leads to positive selection of bacteria harboring resistance determinants and it's spread by horizontal gene transfer between different microbes further complicates the situation. Such events of determinant transfer were detected in patients after antibacterial therapy (Crémet et al., 2012, Karami et al., 2007) and were confirmed experimentally in animals and healthy donors (Lester et al., 2004, 2006). Cases of pathogens spread that received resistant genes from the intestinal microbiota have been reported (Crémet et al., 2012; Goren et al., 2010). Thus, the intestinal microbiota is an important reservoir of AR genes, open to infectious agents of socially significant diseases (Jernberg, Cecilia, et al. 2010). At the moment, various approaches have been applied to study and monitor antibiotics resistance using molecular biology, biochemistry and high-throughput sequencing techniques. Modeling approaches to the epidemiological process have also been developed. However, there is a need in studying the capabilities of the mathematical apparatus for predicting and simulating the spread of AR under the spread of infectious agents in the population, which can substantially supplement and expand the possibilities of existing research methods.

Different approaches have been developed to predict and simulate the resistance of individual infectious agents / microbiota. The most frequent approaches to epidemiological simulation is differential equations based approach. However, as noted above, to study a complex disease behaviour and to develop a strategy for intervention, agent based models are more appropriate.

We created a model for the spread of an infection caused by a pathogen, taking into account the possible gain of resistance determinants. In our model there are two states of pathogens: sensitive to treatment and resistant. Our agents could be in several states: healthy; in the incubation period; taking AB therapy at home or in the hospital and being treated in the hospital for another reason, not infection. Changes in agents states occurs according to the corresponding probabilities which are calculated dynamically. There is also a variable level of agents microbiota resistance level, which accumulates through AB therapy and consumption of products containing antibiotics. In the home treatment state, it is likely that the patient will choose the wrong treatment (preliminary reducing the dose or cessation of treatment). At each point of time, modeling assesses the degree of resistance of the pathogen in the town on average, which affects the probability of infection with a pathogen in the next step.

During the regression analysis, the following parameters were identified as being control: the pathogen infection potential, the incubation period, the probability of wrong antibiotic treatment, the pathogen susceptibility to receive antibiotic resistance, the permanent level of gut antibiotic resistance, the probability to be hospitalized during incubation period. An increment of incubation period, the pathogen infection potential and the probability of wrong antibiotic treatment leading to growing the number of infected people and the probability to be infected (Fig. 3a, b). Obviously, only the latter can be controlled. The main tool here is public education, the promotion of "responsible self-treatment" (link) and the availability of a well designed recommendations. Our results suggests that the main contribution to the growth of the average human microbiota resistance level is made by the increase in the permanent level of gut antibiotic resistance in the population (Fig.3c). It is formed by consuming subtherapeutic doses with food, water treated with AB, and wrong treatment. Clearly, monitoring the quality of agricultural products and checking of the AB presence will help cope with this problem. In the case of an average pathogen resistance level, the high incubation period and the combination of the AR acceptance pathogen potential and the permanent level of gut antibiotic resistance are evidenced, which leads to increased values of the our model indicators (Fig. 3d). The most important characteristics that contribute most to changes in disease spread are the pathogen infection potential, incubation period, probability of wrong antibiotic treatment. These parameters affect all four indicators of the model (the number of infected people, the probability to be infected, the average microbiota resistance level and of the pathogen separately). The effect of the incubation period and the infection potential seems completely logical and does not contradict modern data (Horn, et al., 2013, Leclerc, Melen, et al., 2014; Salje, Henrik, et al., 2016). In general, there are two points of view regarding the dose of the antibacterial drug taken. The first consider that stopping the course of an antibiotic is a safe and effective way to avoid excessive use of antibiotics in many situations (Horne, Rob, et al., 2013, Llewelyn, Martin J., et al, 2017). Others - it is necessary to take AB all the prescribed course (Bassetti, et al., 2015, Tepekule, et al., 2017). The study of the VERA model showed that the probability of wrong treatment (non-compliance with the prescribed course of taking AB) positively affects the indicators. The microbiota resistance and pathogen resistance increase is also influenced by the permanent level of antibiotic resistance of the agent's gut microbiota (accumulated through consumption of food treated with AB) (Moore et al., 2013; Founou et al., 2016; Hoang et al., 2017). Thus, based on our model, it can be concluded that the excessive consumption of antibacterial drugs (here, improper therapy and consumption of food processed by AB) leads to the accumulation of AP microbiota, the spread of AR and to the complicated control of the epidemiological situation of this infection.

According to the analysis of the relationship of indicators among themselves, the number of infected agents, comparable to a pandemic, depends exponentially on the high proportion of resistant pathogens (Fig. 5, b). This could be very important because of catastrophic effects of even small increase in the average pathogen resistance. Since the resistance of pathogens could be investigated, it is important to constantly monitor AR. It is desirable to put into practice the control of the resistance of microorganisms in the main public places of the town. Such information could change the treatment protocols for pathogenic diseases well-timed and not allow catastrophic consequences.

We studied the model using the example of shigellosis. The choice of shigellosis is dictated by the ease of distribution in children's institutions, in places of public catering, in hospitals, etc. [Ref]. CDC in its report [Ref] notes 7,638 cases of infection in the United States in 2013. In 2014 the incidence rate is 6.5 cases per 100,000 people [Ref]. The second, *Shigella spp.* are very prone to antibiotic resistance gain, which can add complications to patients [Ref]. In 2017, WHO published a report, which included Shigella spp. in the list of pathogens that needs development of new AB [Ref]. It is known that shigellosis is most active in summer and autumn, during holidays and travel. Namely, after massive international travel, it is believed that drug resistance outbreaks appear in the United States [Ref, 1].

In our model there are a number of limitations. We do not consider a lethal outcome. The hospital's capacity is equal to the population size. AR is acquired only through horizontal genes transfer. Treatment in the hospital do not leads to AR acquirements, moreover it decreases to some constant value. Also in our model seasonality is not taken into account. However, this may be an advantage, because it could lower the noise associated with additional factors, like the time of year. With the development of NGS technology and the generation of an increasing number of metagenomic samples, it is possible to improve the accuracy of the model, modify it to produce close to more accurate results.

An important characteristic of the model is the combination of factors of possible improper treatment and the formation of a constant level of AR intestinal microbiota. The advantage of this model lies in the ease of obtaining data on the development of AR and the pandemic of the bacterial pathogen with a varying input parameters. Such real world data is often not available. Simulations using model like VERA could be useful in predicting the development of the epidemiological situation associated with the spread of infection with the pathogen. The functional for assessing the optimal observation intensity could also be effective for choosing optimal duration of quarantine, for example, in children's institutions. For illustrative purposes we also implemented a visualization tool which shows model behaviour dynamically and as a pdf-report.

## References

O’Neill J. (2016). Tackling drug-resistant infections globally: Final report and recommendations The review on antimicrobial resistance. [https://amr-review.org]

Rolain J.-M. (2013) Food and human gut as reservoirs of transferable antibiotic resistance encoding genes. Front. Microbiol., 4.

Rhouma, M., Beaudry F., and Letellier A. “Resistance to colistin: what is the fate for this antibiotic in pig production?.” International journal of antimicrobial agents 48.2 (2016): 119–126.

Liu, Yi-Yun, et al. “Emergence of plasmid-mediated colistin resistance mechanism MCR-1 in animals and human beings in China: a microbiological and molecular biological study.” The Lancet infectious diseases 16.2 (2016): 161–168.

Nguyen, Nhung T., et al. “Use of colistin and other critical antimicrobials on pig and chicken farms in southern Vietnam and its association with resistance in commensal Escherichia coli bacteria.” Applied and environmental microbiology 82.13 (2016): 3727–3735.

Österberg, Julia, et al. “Antibiotic resistance in Escherichia coli from pigs in organic and conventional farming in four European countries.” PloS one 11.6 (2016): e0157049.

Sommer M.O. et al. (2009) “Functional characterization of the antibiotic resistance reservoir in the human microflora”. Science, 325, 1128–1131.

Fofana, Mariam O., et al. “A multistrain mathematical model to investigate the role of pyrazinamide in the emergence of extensively drug-resistant tuberculosis.” Antimicrobial agents and chemotherapy 61.3 (2017): e00498–16.

Weinstein, Robert A., et al. “Understanding the spread of antibiotic resistant pathogens in hospitals: mathematical models as tools for control.” Clinical Infectious Diseases 33.10 (2001): 1739–1746.

D’Agata, Erika MC, et al. “Modeling antibiotic resistance in hospitals: the impact of minimizing treatment duration.” Journal of theoretical biology 249.3 (2007): 487–499.

Bergstrom, Carl T., Monique Lo, and Marc Lipsitch. “Ecological theory suggests that antimicrobial cycling will not reduce antimicrobial resistance in hospitals.” Proceedings of the National Academy of Sciences of the United States of America 101.36 (2004): 13285–13290.

Webb, Glenn F., et al. “A model of antibiotic-resistant bacterial epidemics in hospitals.” Proceedings of the National Academy of Sciences of the United States of America 102.37 (2005): 13343–13348.

Austin, Daren J., et al. “Vancomycin-resistant enterococci in intensive-care hospital settings: transmission dynamics, persistence, and the impact of infection control programs.” Proceedings of the National Academy of Sciences 96.12 (1999): 6908–6913.

Cohen, Ted, and Megan Murray. “Modeling epidemics of multidrug-resistant M. tuberculosis of heterogeneous fitness.” Nature medicine 10.10 (2004): 1117–1121.

von Wintersdorff, Christian JH, et al. “Dissemination of antimicrobial resistance in microbial ecosystems through horizontal gene transfer.” Frontiers in microbiology 7 (2016): 173.

Wu, Guojun, et al. “Diminution of the gut resistome after a gut microbiota-targeted dietary intervention in obese children.” Scientific reports 6 (2016).

Yarygin K.S. et al. (2017) Resistomap—online visualization of human gut microbiota antibiotic resistome. Bioinformatics, 33, 2205–2206.

Pehrsson, Erica C., et al. “Interconnected microbiomes and resistomes in low-income human habitats.” Nature 533.7602 (2016): 212.

Hoang, Phuong Hoai, et al. “Antimicrobial resistance profiles and molecular characterization of Escherichia coli strains isolated from healthy adults in Ho Chi Minh City, Vietnam.” Journal of Veterinary Medical Science 79.3 (2017): 479–485.

Moore, Aimee M., et al. “Pediatric fecal microbiota harbor diverse and novel antibiotic resistance genes.” PloS one 8.11 (2013): e78822.

Leclerc, Melen, et al. “Estimating the delay between host infection and disease (incubation period) and assessing its significance to the epidemiology of plant diseases.” PloS one 9.1 (2014): e86568.

Horn, B. J., and R. J. Lake. “Incubation period for campylobacteriosis and its importance in the estimation of incidence related to travel.” Eurosurveillance 18.40 (2013): 20602.

Salje, Henrik, et al. “How social structures, space, and behaviors shape the spread of infectious diseases using chikungunya as a case study.” Proceedings of the National Academy of Sciences 113.47 (2016): 13420–13425.

Bassetti, Matteo, et al. “Preventive and therapeutic strategies in critically ill patients with highly resistant bacteria.” Intensive care medicine 41.5 (2015): 776–795.

Tepekule, Burcu, et al. “Modeling antibiotic treatment in hospitals: A systematic approach shows benefits of combination therapy over cycling, mixing, and mono-drug therapies.” PLoS computational biology 13.9 (2017): e1005745.

Jernberg, Cecilia, et al. “Long-term impacts of antibiotic exposure on the human intestinal microbiota.” Microbiology 156.11 (2010): 3216–3223.

Llewelyn, Martin J., et al. “The antibiotic course has had its day.” Bmj 358 (2017): j3418.

Horne, Rob, et al. “Understanding patients’ adherence-related beliefs about medicines prescribed for long-term conditions: a meta-analytic review of the Necessity-Concerns Framework.” PloS one 8.12 (2013): e80633.

Founou, Luria Leslie, Raspail Carrel Founou, and Sabiha Yusuf Essack. “Antibiotic resistance in the food chain: a developing country-perspective.” Frontiers in microbiology 7 (2016): 1881.

Willem, Lander, et al. “Optimizing agent-based transmission models for infectious diseases.” BMC bioinformatics 16.1 (2015): 183.

Eubank, Stephen, et al. “Modelling disease outbreaks in realistic urban social networks.” Nature 429.6988 (2004): 180.

Colizza, Vittoria, et al. “Modeling the worldwide spread of pandemic influenza: baseline case and containment interventions.” PLoS medicine 4.1 (2007): e13.

Kaufman, James, Stefan Edlund, and Judith Douglas. “Infectious disease modeling: Creating a community to respond to biological threats.” Statistical Communications in Infectious Diseases 1.1 (2009).

Lipsitch, Marc, Carl T. Bergstrom, and Bruce R. Levin. “The epidemiology of antibiotic resistance in hospitals: paradoxes and prescriptions.” Proceedings of the National Academy of Sciences 97.4 (2000): 1938–1943.

Spicknall, Ian H., et al. “A modeling framework for the evolution and spread of antibiotic resistance: literature review and model categorization.” American journal of epidemiology 178.4 (2013): 508–520.

Pitman, Richard, et al. “Dynamic transmission modeling: a report of the ISPOR-SMDM Modeling Good Research Practices Task Force Working Group-5.” Medical decision making 32.5 (2012): 712–721.

Van Bunnik, B. A. D., and M. E. J. Woolhouse. “Modelling the impact of curtailing antibiotic usage in food animals on antibiotic resistance in humans.” Royal Society open science 4.4 (2017): 161067.

D’Costa, Vanessa M., et al. “Antibiotic resistance is ancient.” Nature 477.7365 (2011): 457.

Perron, Gabriel G., et al. “Functional characterization of bacteria isolated from ancient arctic soil exposes diverse resistance mechanisms to modern antibiotics.” PLoS One 10.3 (2015): e0069533.

Levy, Stuart B., and Bonnie Marshall. “Antibacterial resistance worldwide: causes, challenges and responses.” Nature medicine 10.12s (2004): S122.

